# Upper respiratory tract bacterial microbiome features associated with *Streptococcus pneumoniae* colonisation and environmental factors among healthy older children in a rural setting in The Gambia

**DOI:** 10.64898/2026.10.04.756419

**Authors:** Dam Khan, Brenda Kwambana-Adams, Peggy-Estelle Tientcheu, Shola-Able Thomas, Josh L. Espinoza, Gavin John, Tiana Mamaghani, Peter Sylvanus Ndow, Ma-Ansu Kinteh, Archibald Worwui, Isaac Darko Otchere, Nuredin Ibrahim Mohammed, Mark P. Nicol, Chris L. Dupont, Martin Antonio

**Affiliations:** Medical Research Council Unit The Gambia at the London School of Hygiene & Tropical Medicine, Banjul, The Gambia; Department of Infection Biology, Faculty of Infectious and Tropical Diseases, London School of Hygiene & Tropical Medicine, London, United Kingdom; Malawi Liverpool Wellcome Programme, Blantyre, Malawi; NIHR Global Health Group on Vaccines to Control Respiratory Pathogens and AMR, Division of Infection and Immunity, University College London, London, United Kingdom; J. Craig Venter Institute, La Jolla, CA, United States of America; Department of Bacteriology, Noguchi Memorial Institute for Medical Research, Legon, University of Ghana, Accra, Ghana; Marshall Centre, Division of Infection and Immunity, School of Biomedical Sciences, University of Western Australia, Perth, Australia; Centre for Epidemic Preparedness and Response, London School of Hygiene & Tropical Medicine, London, United Kingdom

**Keywords:** *Streptococcus pneumoniae*, upper respiratory tract infections, microbiome, older children

## Abstract

The upper respiratory tract (URT) microbiome plays a vital role in respiratory health, with older children (5-14 years old) serving as key reservoirs for the transmission of bacteria to younger individuals (< 5 years old). However, the URT microbiome in this age group remains poorly characterised, particularly in relation to colonisation by the pathobiont *Streptococcus pneumoniae*. To address this gap, we analysed URT samples collected over eight visits from 134 healthy children aged 5-14 years in The Gambia, using culture-dependent methods to detect *S. pneumoniae* and culture-independent 16S rRNA sequencing to characterise the microbiome in parallel. 16S rRNA sequencing identified *Streptococcus* and *Prevotella* as the dominant microbiome constituents. When integrating longitudinal sampling data, we observed temporal instability with frequent switches in microbiome profiles over time. URT samples colonised with *S. pneumoniae* were associated with increased abundance of *Moraxella*, *Streptococcus* and *Haemophilus,* whereas non-colonised URT samples were enriched with *Granulicatella* and *Filifactor*. *S. pneumoniae* colonisation was also associated with lower Shannon diversity compared with non-colonised samples. Age and season were the strongest determinants of microbiome composition; increasing age was associated with decreasing prevalence and abundance of taxa including *Haemophilus* and *Moraxella*, while gram-positive taxa were predominantly enriched in the dry season. These findings provide new insight into the URT microbiome of older children in The Gambia and highlight *Granulicatella* and *Filifactor* as potential contributors to colonisation resistance against *S. pneumoniae*.

## Introduction

The upper respiratory tract (URT) harbours a complex community of microorganisms whose interactions with each other and with the host immune system play an important role in respiratory health (1). Among the many microbial inhabitants of the URT is the pathobiont *Streptococcus pneumoniae,* a gram-positive diplococcus that asymptomatically colonises the mucosal surfaces of the URT of humans (2). Despite often existing as a harmless commensal, it remains a major cause of severe infections including meningitis, pneumonia, and sepsis in children and adults. In 2021, *S. pneumoniae* was responsible for the highest proportions of lower respiratory tract infection worldwide, with an estimated 97·9 million (92·1-104·0) episodes and 505, 000 deaths (454, 000–555, 000) (3). *S. pneumoniae* colonisation in the URT provides a reservoir for transmission, marking the first step in invasive disease pathogenesis in susceptible individuals. Progression from colonisation to invasive disease may result from URT microbial community imbalance, expression of virulence genes and damage to the host respiratory epithelium (4). The observed substantial variability in pneumococcal disease rates by age and geography mirrors the variation seen in carriage which further highlights the important role of colonisation in disease (5).

Studies of *S. pneumoniae* interactions with other bacteria are not uncommon, although they are commonly in children under five years old, and the focus is gradually shifting from targeted approaches to methods that characterise the repertoire of bacteria in the URT, such as 16S rRNA sequencing and shotgun metagenomics. However, most of these studies remain cross-sectional. Cross-sectional designs are limited because they do not account for seasonal variation which markedly influences microbiome community structure.

More broadly, epidemiological studies investigating the structure of the URT microbiome among older children (5-14 years old) in African and non-African populations are scarce, as noted by these systematic reviews (6, 7). Older children are essential to consider, as they play a crucial role in gatekeeping respiratory health by serving as reservoirs for introducing and transmitting respiratory bacteria to young children, who experience the greatest burden of respiratory illnesses. In particular, *S. pneumoniae* colonisation in older children has been identified as one of the main drivers of PCV13 vaccine-type pneumococcal transmission (8). Moreover, older children accounted for up to 25% of meningitis cases during a pneumococcal meningitis outbreak reported in Ghana in the past decade (9).

We conducted this longitudinal study to examine the association between URT microbiome composition and *S. pneumoniae* colonisation in healthy, older children in a rural Gambian population. Given that the URT is directly exposed to the external environment and its composition shifts in response to environmental factors, we also described the environmental and sociodemographic factors shaping URT microbiome composition in this setting.

## Methodology

### Study design, participants and specimen collection

Sample collection for this longitudinal, observational, prospective study was conducted from February 2019 to March 2020, and from February 2021 to March 2022. Field activities were suspended between April 2020 to January 2021 due to COVID-19 pandemic mitigation measures. Sampling occurred every three months in four visits before the suspension period and four visits after the suspension period making a total of eight visits. This study was conducted in Foni Bintang-Karenai, located in the West Coast Region of The Gambia. Face-to-face surveys were conducted to obtain the health and demographic details of the participants. Exclusion criteria included participants with pre-existing medical conditions for which they were receiving antibiotic treatment, and those who permanently relocated outside the study area during follow-up. Combined nasopharyngeal and oropharyngeal (NP/OP) specimens were collected by trained nurses using FLOQSwabs® (Copan Diagnostics) at each visit. Data were also collected on environmental and sociodemographic covariates using the REDCap software on electronic tablets. All collected NP/OP specimens were immediately placed in Skim milk-Tryptone-Glucose-Glycerol medium (STGG) and transported 85 km on ice to the Biobank in batches, where they were stored at −70°C until processing. A summary of the study methodology is illustrated in Figure 1.

**Figure 1:**
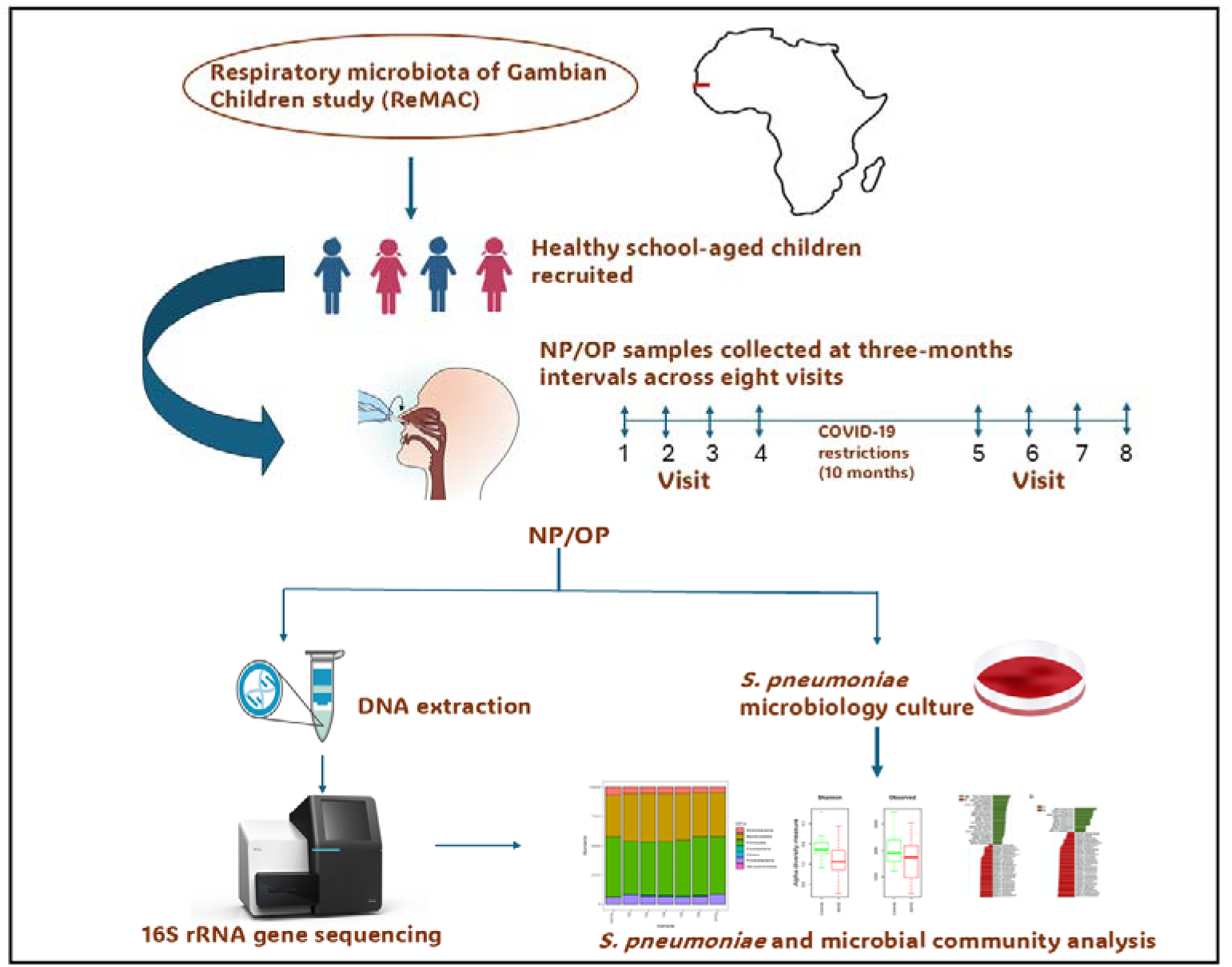
Summary of study workflow

## Microbiology culture and DNA extraction

Up to 10 µl of thawed NP/OP was inoculated onto Columbia agar supplemented with 5% (v/v) defibrinated sheep blood and 5 g/L of gentamicin using the quadrant streaking method. The plates were incubated at 37°C in 5% CO2 overnight (16-24h). Presumptive *S. pneumoniae* were identified based on alpha haemolysis and colony morphology and subcultured on Columbia blood agar. Optochin susceptibility was determined, and samples exhibiting a zone of inhibition of ≥ 14 mm were identified as *S. pneumoniae.* DNA extraction was performed on the same NP/OP samples using the QIAGEN DNeasy PowerLyzer PowerSoil kit. Briefly, 200 µl of NP/OP specimens were added to extraction beads and SDS-containing solution and vortexed for 15 minutes to aid in cell lysis. The supernatant was sequentially treated with two Inhibitor Removal Technology solutions® to precipitate inhibitors, such as cell debris and proteins. The purified supernatant was mixed with a salt-binding solution and passed through a silica spin column for DNA binding. After ethanol-based washing, the DNA was eluted in 100 µl elution buffer.

The Microbiology culture and DNA extraction was carried out at the WHO Regional Reference Laboratory host by the Medical Research Council unit The Gambia at the London School of Hygiene and Tropical Medicine. The Laboratory participates in external quality assurance (EQA) programmes organised by the United Kingdom National External Quality Assessment Service (UK NEQAS), the National Institute for Communicable Diseases (NICD), South Africa, and the United States Centers for Disease Control and Prevention (US CDC).

### 16S ribosomal RNA gene sequencing

16S ribosomal RNA gene amplification was performed on the extracted NP/OP samples at the J. Craig Venter Institute, San Diego using Forward primer 515F-Y (5’-GTGYCAGCMGCCGCGGTAA) and Reverse primer 926R (5’-CCGYCAATTYMTTTRAGTTT), which span the V4/V5 hypervariable regions of the 16S rRNA gene. The forward and reverse primers were combined in 96 well plates using an automated liquid handling robot (Eppendorf epMotion® 5075). The total volume of the PCR amplification step was 25 µl, and each reaction contained 10 µM of 1 µl combined forward and reverse primer mix, 5 µl of 5X Azura buffer, 0.25 µl of Trufi DNA polymerase, 16.75 µl of molecular-grade water, and 2 µl of DNA template. The Thermal cycling conditions were 95°C for 1 minute, followed by 30 cycles at 95°C for 15 seconds, 57°C for 30 seconds, and an extension of 72°C for 1 minute. PCR products were cleaned and normalised using the SequalPrep™ kit protocol (Invitrogen, Catalog # A10510-01), after which 20 µl of each sample was pooled. The pooled libraries were analysed on an Agilent 2100 Bioanalyzer using a high sensitivity DNA chip (Agilent, part No. 5067-4626). DNA quantification of the final library was performed using the High Sensitivity dsDNA Qubit assay (ThermoFisher, catalogue number Q32854). The final library with 10% PhiX was sequenced using a MiSeq Illumina sequencer. Blank field controls, negative extraction controls and PCR controls were also amplified and sequenced in parallel with the samples.

### 16S rRNA sequence data processing

Raw paired-end read processing, including quality control, denoising, chimera removal, and trimming was performed using DADA2 v1.26.0 (10). The processed sequencing reads were dereplicated and Amplicon Sequence Variants (ASVs) inferred. Taxonomic classification of each ASV was assigned using the SILVA v138.1 database (11). The DADA2 outputs were converted to tab-separated value (TSV) and FASTA files for use in all abundance and ecological analysis. ASVs classified as Bacteria were retained while those classified as Eukaryota, Archaea, Chloroplast at the order level, and those with no taxonomic assignment at the phylum level were excluded from the dataset. We used the prevalence method of the decontam (12) R package (threshold = 0.10) to identify and remove contaminant ASVs derived from sampling, DNA extraction or PCR reagents. Based on this method, 23 ASVs, of which 43% (10/23) were *Cutibacterium* were removed. Samples with fewer than 1000 reads after contaminant removal were also removed from the dataset, in line with common practice. This threshold also ensured that the retained samples had read counts distinct from those of the negative controls (Supplementary Information, Figure1). ASVs that were not represented at a minimum of 0.1% relative abundance in at least 1% of the samples were excluded, as these might be artifacts or taxa that are too uncommon to have an impact on the outcome measurements.

### Statistical analyses

RStudio software v.4.4.2 was used for data analysis and visualization. Bar plots of the mean compositional relative abundance at the genus and phylum levels were plotted using ggplot2 v4.0.3. We assigned each sample to a microbiome profile defined by the genus with the highest relative abundance in the sample. Transitions in microbiome profiles of each sample were visualised using Alluvial plots (ggalluvial v0.12.5) (13). We used a mixed-effects logistic regression model to estimate the association between *S. pneumoniae* colonisation with season and age. A linear mixed-effects model implemented in the lme4 v1.1.37 package was used to evaluate the association between *S. pneumoniae* colonisation and Shannon diversity, adjusting for age at visit and season of sample collection with *ParticipantID* as random effect. We computed Aitchison distance for the beta diversity analysis of the microbiome, stratified by *S. pneumoniae* colonisation state, and principal coordinate analysis (PCoA) plots was used for visualisation. Up to 1000 permutations of the PERMANOVA test was applied using the *adonis2* function of the vegan v2.6.8 package to determine differences in beta diversity. *ParticipantID* was set as a blocking variable. A P-value of <0.05 was considered significant in the Shannon and beta diversity analysis. We used Microbiome Multivariable Associations with Linear Models (MaAsLin3 v0.99.2) to test for differences in abundance and prevalence of bacteria at the genus level in *S. pneumoniae* colonised and non-colonised states. We applied a Centered Log-Ratio (CLR) transformation of the abundance data in the model and set *ParticipantID* as random effect. Sequencing read depth was included as a covariate to control for prevalence associations owing to uneven sequencing depths. An association of q < 0.1 (MaAsLin3 default) was considered significant after correcting for multiple testing using the Benjamini-Hochberg correction. Joint q-value as indicated in the MaAsLin3 output is reported for taxa showing significant associations in both the prevalence and abundance models. For taxa significant in only one model, the q-value from the respective model is reported individually. Season and age at sample collection were controlled for in the models. We also accounted for repeated sampling and potentially correlated observations from the same individual in the models.

Shannon diversity and MaAsLin3 analysis was also performed for other covariates to study the impact of environmental and sociodemographic factors on microbiome composition. Other covariates tested included season, age at sample collection, fuel source for cooking, waste disposal method, number of siblings, presence of pets (cats and/or dogs) in the compound, and presence of livestock (cattle, sheep, and goats) in the compound.

## Results

### Samples analysed and participant characteristics

A total of 885 NP/OP of the 1,072 expected (82.6%) samples were collected from 134 participants across eight visits. Participant withdrawals from the study were mainly due to relocation from the study area (data not shown). Microbiology culture and 16S rRNA sequencing were performed for all the 885 samples. Microbiology culture results were available for all the samples for analysis. 16S rRNA sequencing data was available for 799/885 (90.3%) samples for analysis following quality control processes. The number of samples collected at each visit and those retained for microbiome analysis following quality control processes are shown in Supplementary Information, specifically Table S1. At least 82.8% of the samples were retained at each visit. Supplementary Information, Table S2 and Figure S2 show how many of the 134 participants contributed between 0 and 8 samples. A total of 42/134 (31.3%) participants had their samples analysed across all eight visits. The baseline (visit 1) environmental and sociodemographic characteristics of the participants are shown in Table 1. Of the 134 children, 50.7% (68/134) were female and 49.3% (66/134) were male. Most participants at baseline were aged 5-7 years, 52.2% (70/134).

**Table 1:**
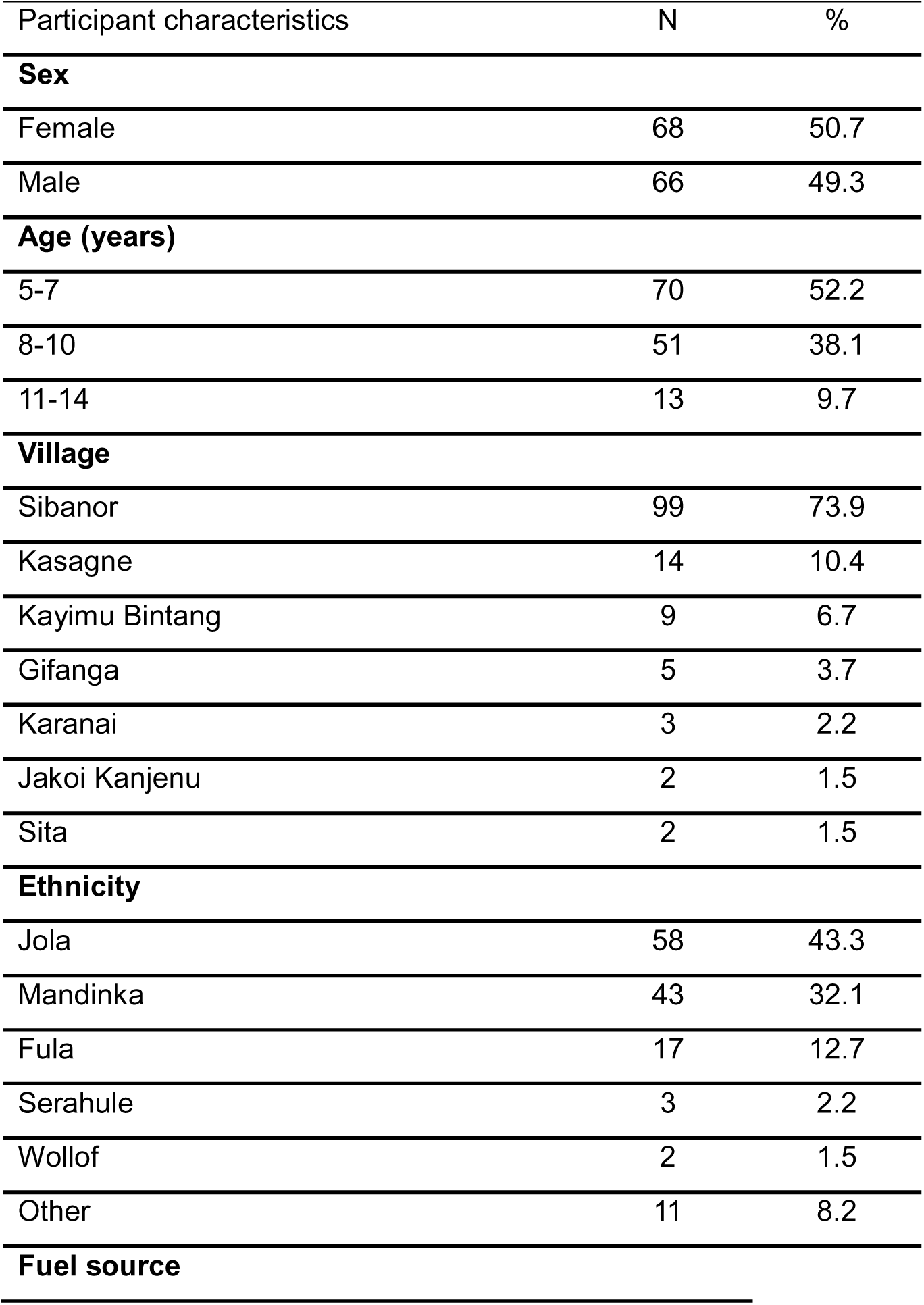
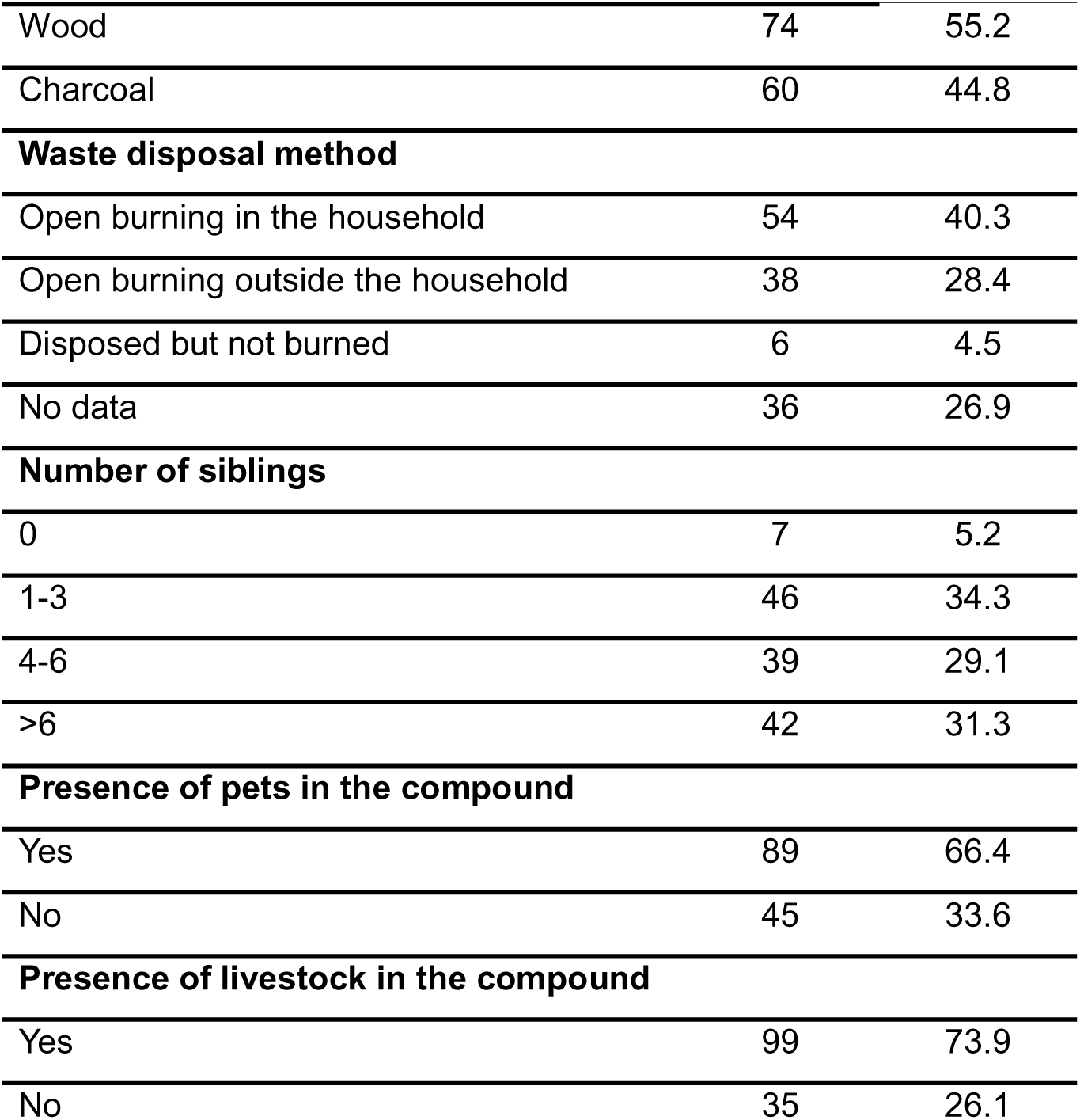
Characteristics of study participants at baseline (Visit 1)

| Participant characteristics | N | % |
| --- | --- | --- |
| **Sex** |  |  |
| Female | 68 | 50.7 |
| Male | 66 | 49.3 |
| **Age (years)** |  |  |
| 5-7 | 70 | 52.2 |
| 8-10 | 51 | 38.1 |
| 11-14 | 13 | 9.7 |
| **Village** |  |  |
| Sibanor | 99 | 73.9 |
| Kasagne | 14 | 10.4 |
| Kayimu Bintang | 9 | 6.7 |
| Gifanga | 5 | 3.7 |
| Karanai | 3 | 2.2 |
| Jakoi Kanjenu | 2 | 1.5 |
| Sita | 2 | 1.5 |
| **Ethnicity** |  |  |
| Jola | 58 | 43.3 |
| Mandinka | 43 | 32.1 |
| Fula | 17 | 12.7 |
| Serahule | 3 | 2.2 |
| Wollof | 2 | 1.5 |
| Other | 11 | 8.2 |
| **Fuel source** |  |  |
| Wood | 74 | 55.2 |
| Charcoal | 60 | 44.8 |
| **Waste disposal method** |  |  |
| Open burning in the household | 54 | 40.3 |
| Open burning outside the household | 38 | 28.4 |
| Disposed but not burned | 6 | 4.5 |
| No data | 36 | 26.9 |
| **Number of siblings** |  |  |
| 0 | 7 | 5.2 |
| 1-3 | 46 | 34.3 |
| 4-6 | 39 | 29.1 |
| >6 | 42 | 31.3 |
| **Presence of pets in the compound** |  |  |
| Yes | 89 | 66.4 |
| No | 45 | 33.6 |
| **Presence of livestock in the compound** |  |  |
| Yes | 99 | 73.9 |
| No | 35 | 26.1 |

### Taxonomic composition of the URT microbiome

Sequencing reads were classified into 3196 ASVs representing 8 phyla and 69 genus-level groups. The largest proportion of bacteria at the genus level (47.8%, 33/69) belonged to the phylum Firmicutes, while Bacteroidota and Proteobacteria had 17.4% (12/69) and 15.9% (11/69) genus-level groups respectively. Spirochaetota, Patescibacteria and Campylobacterota were each represented by a single genus-level group. *Prevotella* (24.5%) and *Streptococcus* (22.3%) were the most frequently dominant taxa (Figure S3). *Prevotella* and *Alloprevotella* had the highest number of unique ASVs (392 and 247 respectively; Figure S4). The mean relative abundance at the genus and phylum levels by study visit is shown in Figure 2. Across all visits, Firmicutes had the highest mean relative abundance (32.7%), followed by Bacteroidota (26.8%) and Proteobacteria (20.8%). Within Firmicutes, *Streptococcus* had the highest mean relative abundance, with 34.6% of the total abundance of the phylum, followed by *Veillonella* (18.5%) and *Granulicatella* (8.8%). *Prevotella* (39.4%), *Alloprevotella* (27.2%), and *Porphyromonas* (24.1%) had the highest mean relative abundance among the Bacteroidota phylum, while *Neisseria* (27.5%), unclassified *Pasteurellaceae* (19.3%) and *Moraxella* (18.5%) had the highest mean relative abundance in the Proteobacteria phylum. *Streptococcus* (11.3%) and *Prevotella* (10.5%) had the highest mean relative abundance across all genus-level groups, followed by *Alloprevotella* (7.3%).

**Figure 2:**
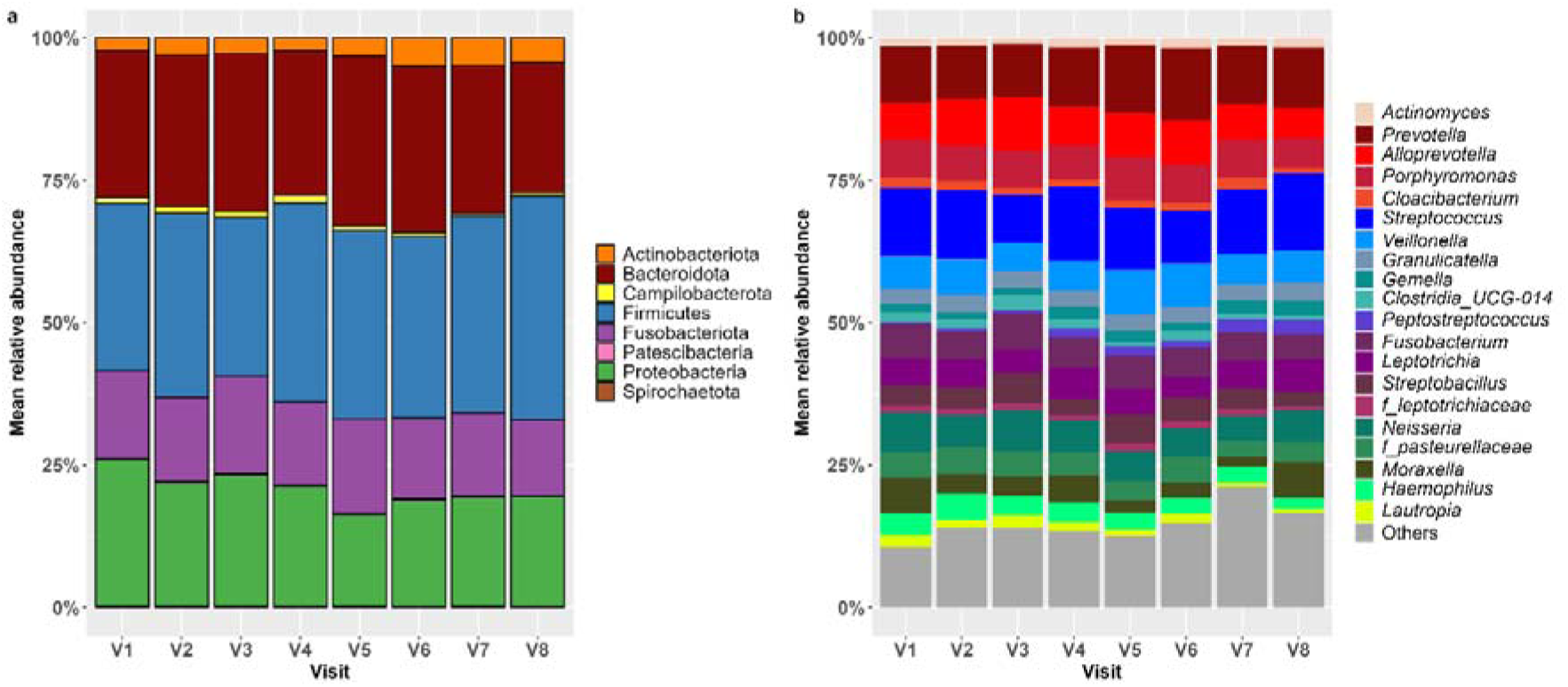
Mean relative abundance of bacterial taxa across eight visits at the phylum (a) and genus (b) level. The phylum-level plot shows all eight phyla in the dataset. The genus-level plot shows the 20 genera with the highest relative abundance while the remaining genera are grouped as “Others”. The genus colour shades in plot b correspond to the colours of their respective phyla in plot a: shades of red represent Bacteroidota, shades of blue represent Firmicutes, shades of purple represent Fusobacteriota, and shades of green represent Proteobacteriota. Genera that are in each phylum are ordered by descending relative abundance.

The median number of bacterial genus-level groups per sample (relative abundance > 0) was 25 (range: 5-43). The URT microbiome was dominated by a small number of taxa; in 86.5% (691/799) of the samples, five or fewer genera accounted for at least 50% of the total relative abundance. We obtained 30 distinct microbiome profiles from the dataset. However, 87.7% (701/799) of the samples were represented by only 10 microbiome profiles. The *Prevotella* microbiome profile was the most common (24.5%, 196/799), followed by *Streptococcus* (22.3%,178/799) and *Alloprevotella* (10.1%, 81/799). We considered a microbiome profile to be stable if the next visit sample from the same individual had the same profile. Microbiome profiles were highly unstable in individuals, with state transitions per visit occurring in 83.2% of the time (Figure 3). Among the profiles represented in at least 10 samples, *Streptococcus* showed the highest retention rate. Individuals maintained the *Streptococcus* profile across consecutive visits 27.3% of the time.

**Figure 3:**
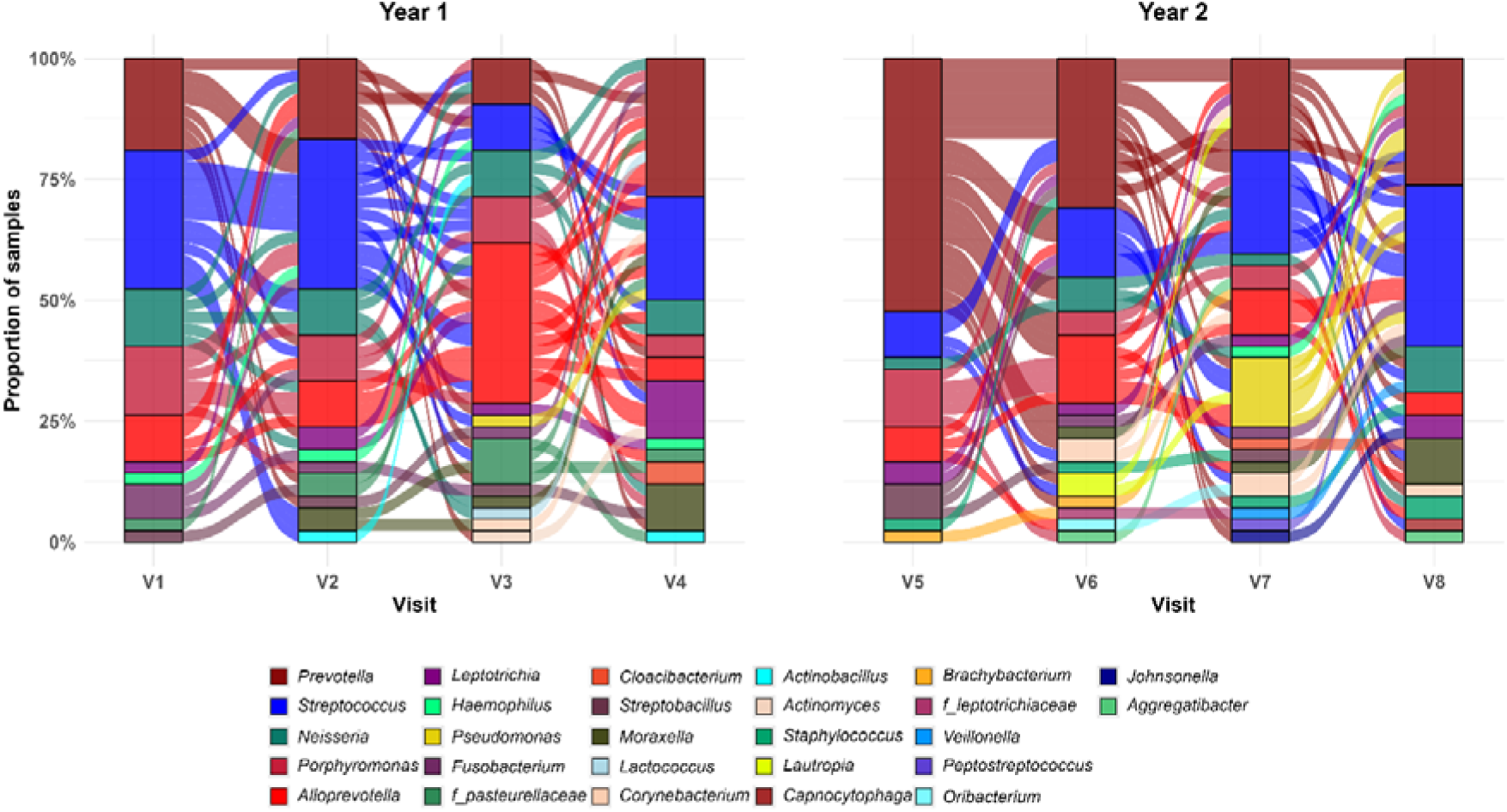
Alluvial plot showing transitions in the upper respiratory microbiome profiles across eight visits among children whose samples were analysed in all eight visits. The y-axis represents the proportion of samples dominated by each microbiome profile. Each flow illustrates transition of dominant microbiome profiles across visits. Data are presented separately for year 1 (V1-V4) and year 2 (V5-V8) due to the ten months gap between visit 4 and 5.

### Culturable *S. pneumoniae* colonisation and the microbiome

URT samples were cultured on selective media to identify individuals colonised or not colonised with *S. pneumoniae*. The overall proportion of *S. pneumoniae* colonisation by culture was 41.2% (365/885). The highest proportion was recorded at Visit 4 (54.4%, 62/114) and the lowest at Visit 7 (23.7%, 23/97). A total of 81 participants had their samples processed by microbiology culture across all eight study visits (Figure 4). The median frequency of *S. pneumoniae* colonisation was three episodes per participant. The highest colonisation frequency was observed in three individuals, each colonised during seven of the eight visits (87.5%). Conversely, two participants were non-colonised in all eight visits. To characterise carriage patterns, participants were categorised by colonisation frequency: High carriers (6–8 episodes), Moderate carriers (3–5 episodes), and Low carriers (0–2 episodes). Most participants were classified as moderate carriers (44/81; 54.3%).

**Figure 4:**
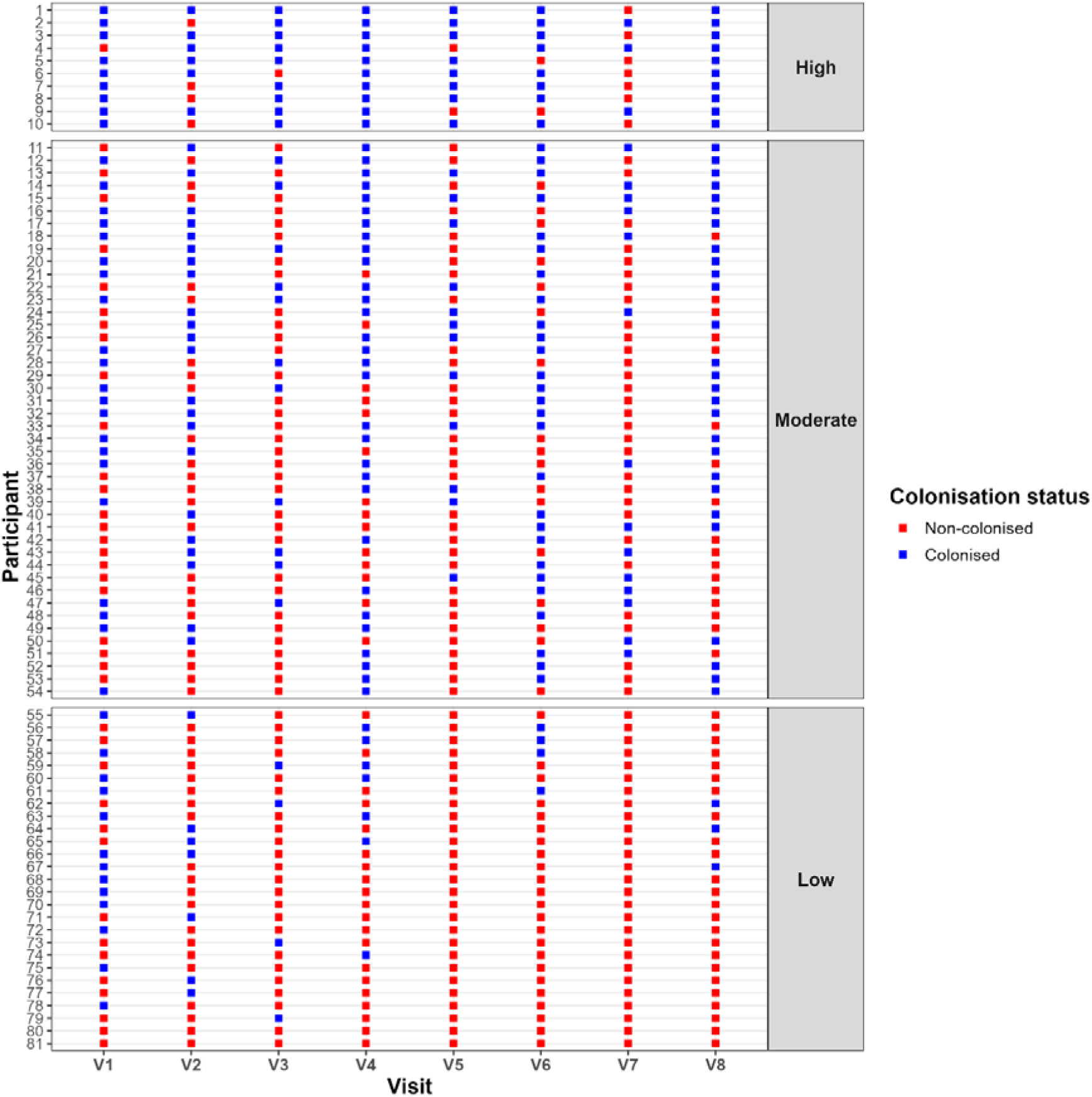
Longitudinal *S. pneumoniae* colonisation profiles of study participants across eight consecutive visits. Trajectories are shown for the 81 participants who completed all eight study visits and had microbiology culture results. Participants are stratified into three groups based on colonisation frequency: High (colonised at 6–8 visits), Moderate (colonised at 3–5 visits), and Low (colonised at 0–2 visits). Each row within a panel represents an individual participant, and each square denotes a single microbiological culture outcome. Blue squares indicate samples from which *S. pneumoniae* was isolated, while red squares indicate samples negative for *S. pneumoniae*.

To assess the persistence of colonisation between consecutive visits, we considered, among the 81 participants with results at all eight visits, every pair of directly consecutive visits in which the participant was colonised at the first visit of the pair. The Visit 4 to Visit 5 interval was excluded from this analysis owing to the extended COVID-19-related gap between those two visits. For each such pair, we determined whether colonisation was maintained at the following visit. Of the 178 consecutive visit-pairs in which the participant was colonised at the first visit, colonisation was maintained at the subsequent visit in 41.0% (73/178) of cases.

The odds of *S. pneumoniae* colonisation decreased with increasing age but this was not statistically significant. For each additional year of age, the odds of colonisation decreased by 5% (OR: 0.95, 95% CI: 0.88–1.03, p=0.204; Figure 5a). The odds of *S. pneumoniae* colonisation were significantly higher in the dry season (November-May) than in the wet season (June-October) (OR: 1.81, 95% CI: 1.34-2.44, p<0.001; figure 5b). Within-sample bacterial diversity (Shannon diversity) was significantly higher in non-colonised samples compared to colonised samples (p<0.001; Figure 5c). Microbiome composition differed between *S. pneumoniae* colonised and non-colonised samples although the variance explained was minimal (R^2^ = 0.45%, p<0.001; Figure 5d).

**Figure 5:**
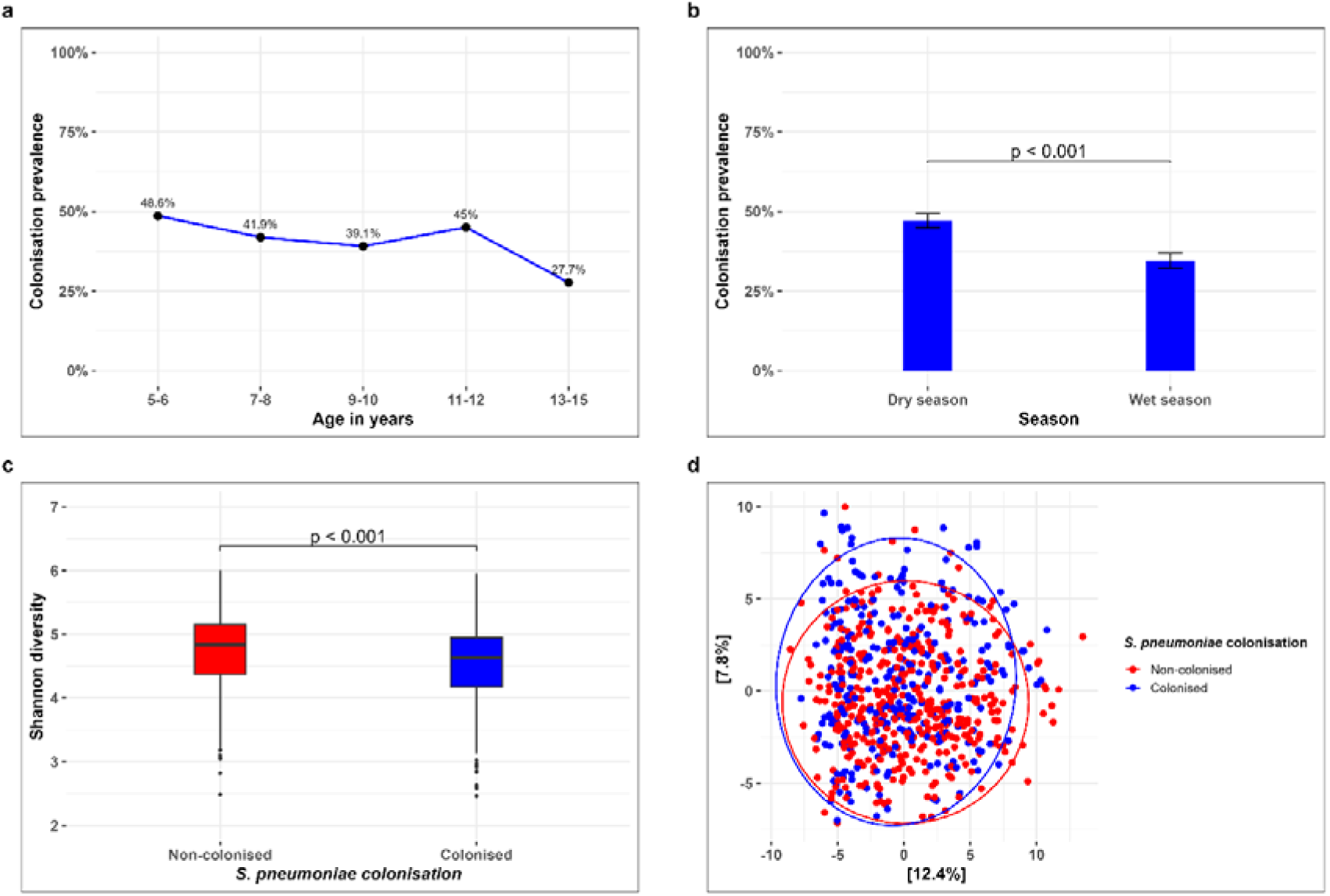
Relationship between *S. pneumoniae* colonisation with age, season and bacterial microbiome diversity. a) Line plot showing colonisation prevalence of *S. pneumoniae* by age at sample collection, with each black dot representing the proportion of colonised children and x-axis showing age range in years. b) Bar plot of *S. pneumoniae* colonisation across the two seasons in The Gambia. The dry season is from November to May yearly while the wet season is from June to October. c) Box plot displaying the Shannon diversity index between samples colonised and non-colonised with *S. pneumoniae*. The horizontal line in the box represents the median index. The bottom edge of the box represents the point below which 25% of the data lies while the top edge of the box represents the point above which 25% of the data lies. The end of the whiskers represents the largest and smallest data values that are not outliers. d) PCoA ordination plot showing the overall differences in microbiome composition between *S. pneumoniae* colonised and non-colonised children based on Aitchison distance.

In our *MaAsLin3* abundance model, higher relative abundance of *Moraxella (q=0.011)*, *Streptococcus (q=0.059)* and *Haemophilus (q=0.059)* were associated with *S. pneumoniae* colonisation (Figure 6), whereas *Filifactor* (q=0.014) relative abundance was higher in *S. pneumoniae* non-colonised samples. Increased relative abundance of *Granulicatella* (q=0.126) was weakly associated with non-colonised samples. In the prevalence associations, *Moraxella* was more frequently detected in *S. pneumoniae* colonised samples (q=0.018), while *Granulicatella* was more common among *S. pneumoniae* non-colonised samples (q=0.035).

**Figure 6:**
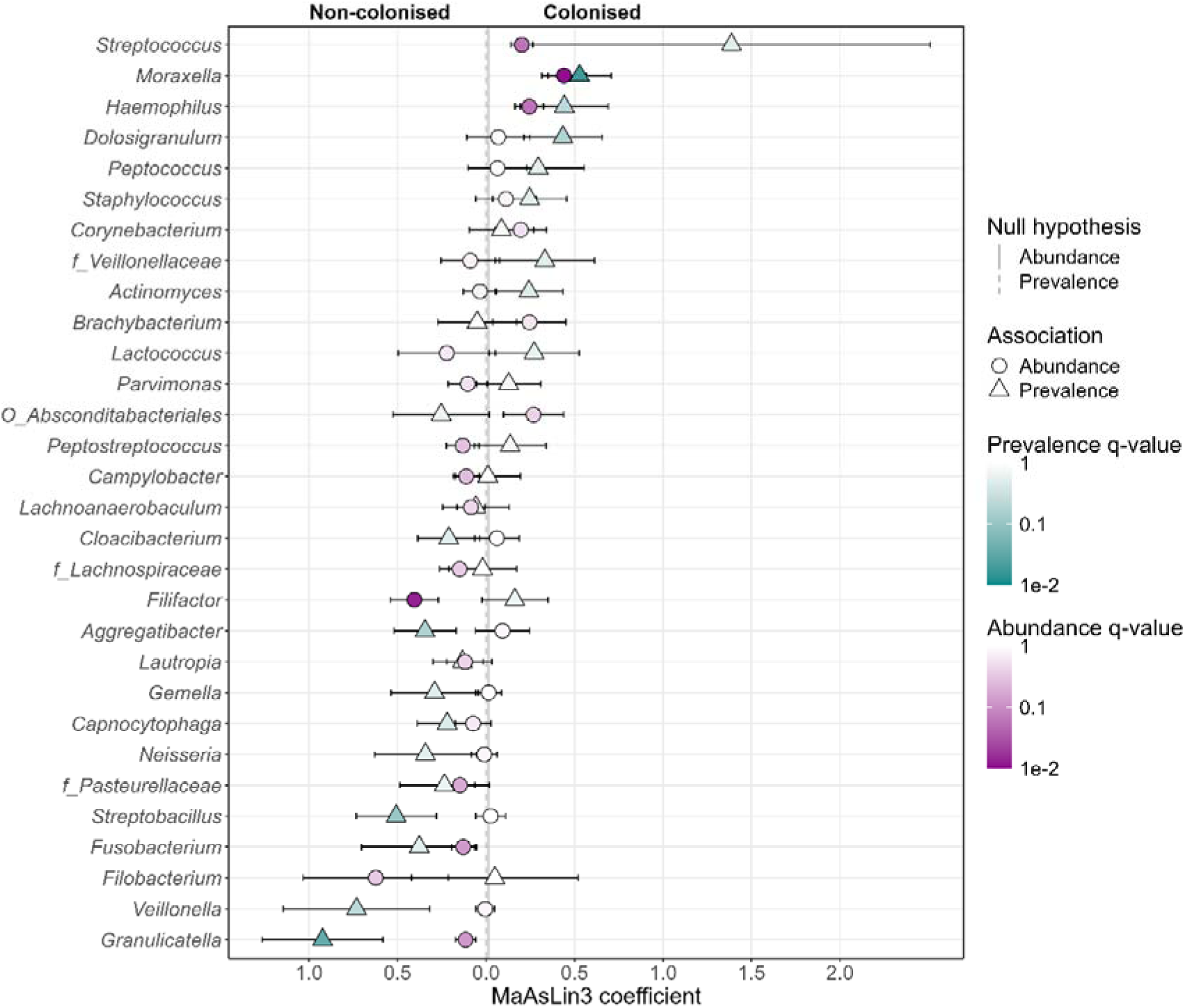
Visualisation of *S. pneumoniae* colonisation MaASLIN3 microbial associations. The circles indicate abundance-based associations and triangles indicate prevalence-based associations. The colour intensity of both shapes represents the adjusted p values, FDR with darker shades representing lower values. The top 30 bacteria with the most important associations are shown on the plot.

### Environmental and demographic factors and the microbiome composition

We evaluated the association between Shannon diversity, relative abundance, and prevalence with environmental and demographic factors. Age at sample collection was the only factor that showed a difference in Shannon diversity analysis with bacterial diversity increasing with age (p= 0.019) (Supplementary Information, Table S2).

In the differential abundance and prevalence analysis, age at sample collection and seasonality showed the strongest association with microbiome composition. Increases in both the prevalence and relative abundance with increasing age were observed in *Peptostreptococcus* (joint q = 2.65 × 10^⁻6^) and *Parvimonas* (joint q = 2.68 × 10^⁻6^), whereas decrease in the prevalence and relative abundance with increasing age was observed in four taxa, namely, *Haemophilus* (joint q = 6.68 × 10^⁻4^)*, Moraxella* (joint q = 2.1 × 10^⁻4^), *Leptotrichia* (joint q = 0.009) and *Campylobacter* (joint q = 0.035) (Figure 7). Of the taxa showing significant seasonal associations, gram-positive taxa predominated in the dry season (6/8, 75%), with increased relative abundance of *Gemella* (q = 0.002), *Streptococcus* (q = 0.004), *Mogibacterium* (q = 0.008), and *Lachnoanaerobaculum* (q = 0.02), and increased prevalence of *Lactococcus* (q = 0.02) and *f_Lachnospiraceae* (q = 0.05). Gram-negative taxa enriched in the dry season included Moraxella (increased prevalence and abundance, joint q = 2.59 × 10^⁻10^) and *Actinobacillus* (increased prevalence, q = 0.05). In the wet season, the distribution between gram-positive (4/7, 57%) and gram-negative taxa (3/7, 43%) was more balanced, with increased prevalence observed in *Brachybacterium* (q = 6.69 × 10^⁻9^), *Staphylococcus* (q = 1.53 × 10^⁻7^), and *f_Staphylococcaceae* (q = 2.3 × 10^⁻4^), and increased prevalence and abundance in *Corynebacterium* (joint q = 5.24 × 10^⁻8^) among gram-positives, and increased prevalence of *Pseudomonas* (q = 1.23 × 10^⁻5^), *O_Absconditabacteriales* (q = 0.001), and *Aggregatibacter* (q = 0.03) among gram-negative taxa

**Figure 7:**
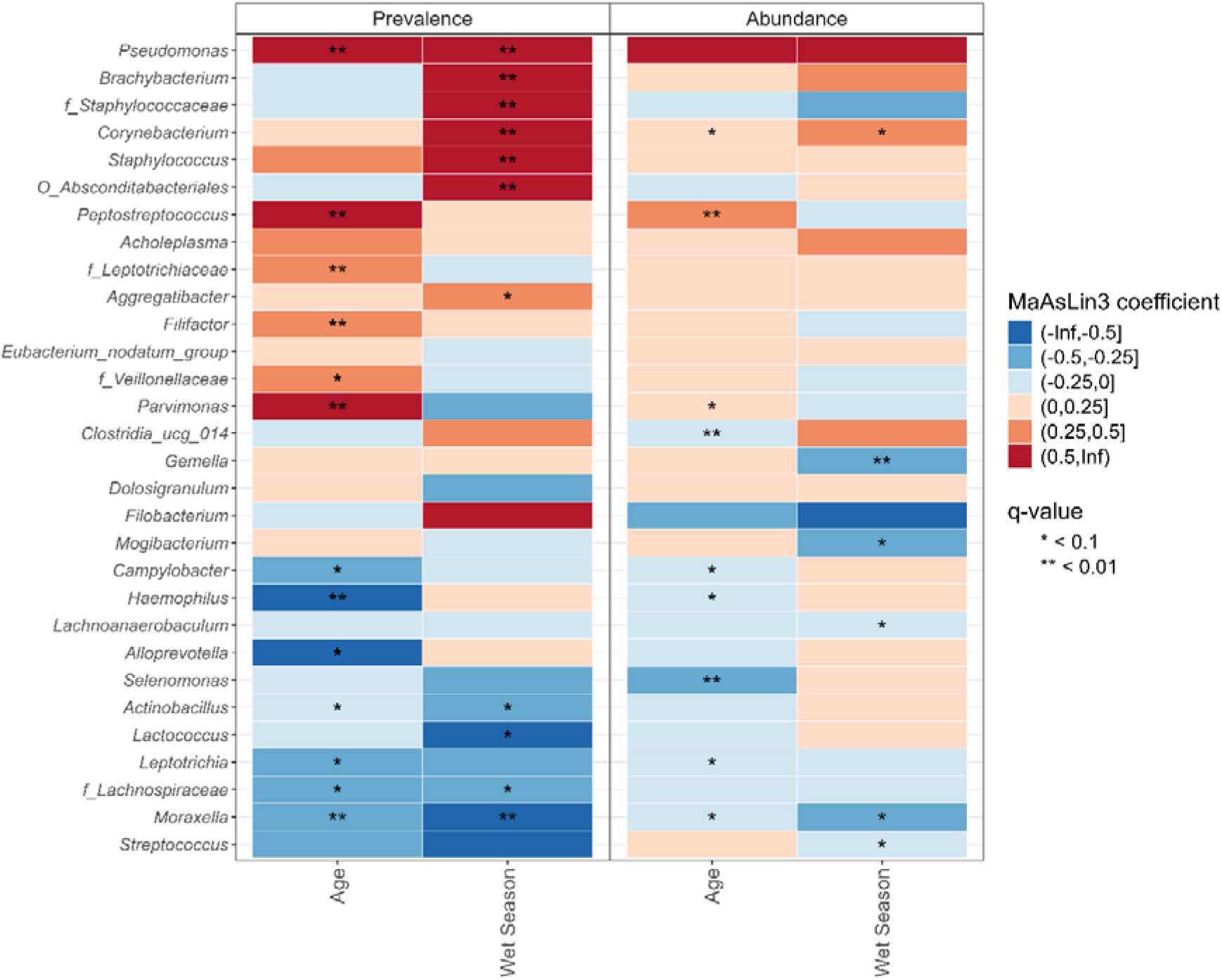
Heatmap of MaASLIN3 microbial associations by age and season. Bacteria that increase in relative abundance or prevalence with increase in age or in wet season are indicated in red shade. Bacteria that increase in relative abundance or prevalence with decrease in age or in dry season are indicated in blue shades. The top 30 bacteria with the most important associations are shown on the plot.

Beyond *S. pneumoniae* colonisation status, season and age, other environmental and sociodemographic factors were associated with URT microbiome composition. Children living in households with livestock had a higher relative abundance of *Campylobacter* (q = 0.063) and higher prevalence of *Lactococcus* (q = 0.049), Supplementary Information, Table S3. Being male was also associated with a higher relative abundance of *Campylobacter* (q = 0.071), Increased abundance of *Megasphaera* (q = 0.042) was associated with living in households with pets. No differences in bacterial composition based on sibling count were observed.

## Discussion

This observational, longitudinal study in rural Gambia characterised the URT microbiome of healthy children aged 5-14 years of age and assessed its association with *S. pneumoniae* colonisation. Our analysis identified *Streptococcus and Prevotella* as the dominant bacteria in the URT. Notably, we observed considerable temporal instability with frequent switch in bacterial profiles over time. In keeping with prior research, age and season emerged as the strongest factors associated with variation in microbiome composition. Shannon diversity was higher in *S. pneumoniae* non-colonised samples and colonisation was lower in those with high prevalence of *Granulicatella* or high relative abundance of *Filifactor*.

While Firmicutes, Bacteroidetes and Proteobacteria dominate the human URT across age groups, representative genera vary considerably (14, 15). Our work addresses a critical gap by focusing specifically on healthy older children in a rural African setting. Here, *Streptococcus and Prevotella* form the backbone of the URT microbiome. A similar study conducted among older children, some of whom had URT infections in Cote D’Ivoire found *Streptococcus and Haemophilus* to be the most common bacteria (16) in the oropharynx. *Prevotella* was the fourth most common bacteria in that cohort. In contrast, research on healthy Dutch children aged 5-14 years reported *Moraxella* and *Corynebacterium* profiles as the most common in the nasopharynx (17).

Early childhood URT communities partially originate from maternal sources (18) and undergo drastic changes in the first month of life before typically stabilising around communities dominated by *Moraxella, Corynebacterium*, *Haemophilus* and *Streptococcus,* as documented in African populations (19–23). Prominence of *Streptococcus* appears to carry through to older children as observed in our study. Despite overall profile instability between visits, nearly one-third of *Streptococcus* dominated profiles persisted in the next timepoint. Persistence of Streptococcus in the upper respiratory tract is facilitated by adhesion to epithelial surfaces through the action of their adhesins and subsequent biofilm formation which promote long term survival in this niche(24).

*Prevotella* is not a common component of the URT in African infants and young children, yet it was very common in our older cohort. Interestingly, high *Prevotella* abundance is a hallmark of the gastrointestinal microbiome of African and other non-westernised populations whose diets are rich in plant-derived fibers and carbohydrates(25–27). The large repertoire of carbohydrate-active enzymes in *Prevotella* enables efficient degradation of plant fibers (28). Supporting this was a study of Gambian toddlers that found *Prevotella copri* and *Prevotella stercorea* comprising up to 42% of the abundance of bacterial species in the gut (29). Our data suggest a potential late ecological acquisition or expansion of *Prevotella* in the URT which is likely influenced by the oropharyngeal component of our samples given that *Prevotella* is common in the oral cavity as it inhabits the gingiva around the teeth (30).

*Moraxella, Haemophilus* and *Campylobacte*r were more common among younger children in our analysis. Enrichment of *Moraxella* in younger participants has also been reported among surgical patients aged 20 weeks to 18 years (31), and in individuals under 21 years with SARS COV 2 infection or exposure (32). In contrast, *Corynebacterium* and *Staphylococcus*, were more common among older children in the studies cited. These two bacteria also appeared more frequently among older children in our study although they were not the most prominent age-associated bacteria. Instead, *Peptostreptococcus* and *Parvimonas* showed the strongest association with older age in our data, even though these taxa are not commonly reported in studies of the upper respiratory tract.

No seasonal differences were observed in Shannon diversity. However, *Corynebacterium* and *Pseudomonas* were more common in the wet season. *Pseudomonas* is widely distributed in environmental habitats, including soil and aquatic environments, and its higher prevalence during the wet season may relate to seasonal changes in environmental conditions, including increased exposure to stagnant water sources following rainfall. *Campylobacter* was overrepresented in children from households with livestock and was more abundant among boys, likely reflecting the gendered patterns of livestock interaction. Many livestock species are known reservoirs for *Campylobacter*. *Campylobacter jejuni* and *Campylobacter coli* are primarily gut pathogens in humans while *Campylobacter concisus* (genomespecies 1) is more frequently detected in oral samples (33). Further characterisation is thus needed to determine which *Campylobacter* species are present in the URT. The high prevalence of *Lactococcus* among children living in households with livestock is more plausibly related to exposure to dairy products and less so on zoonotic transmission. *Lactococci* are abundant in raw milk but are not stable long-term colonisers in the gastrointestinal tracts of cattle limiting their likelihood of sustained animal to human transmission. Close contact with pets may also influence the URT microbiome. *Megasphaera,* which was more abundant in children living in compounds with pets is recognised as a component of the core feline gut microbiome (34)

Our observed *S. pneumoniae* colonisation prevalence (41%) is strikingly similar to estimates from the pre-PCV era. A population-based survey conducted in 2009 among healthy individuals 5-17 years old in rural Gambia reported a prevalence of 42% (35). In addition, a 2009 systematic review on pneumococcal colonisation among children 5–15 years in sub-Saharan Africa also reported a prevalence of 42% (36). This constant colonisation prevalence is linked to serotype replacement where non vaccine serotypes expand to fill the niche left by the vaccine serotypes (37–39). The high rate of *S. pneumoniae* colonisation in the dry season as we have shown is linked to the dry dusty winds experienced during this time of the year (harmattan season) which carry high levels of particulate matter. Environmental exposure to inhaled dust particles has been shown to increase the prevalence of *S. pneumoniae* in the respiratory tract in mouse experimental studies (40). Pneumococcal disease epidemiology studies have also found increase in pneumococcal meningitis in the dry season (41). *S. pneumoniae* colonisation was associated with lower Shannon diversity in our study and it also correlated positively with increased relative abundance of *Moraxella* and *Haemophilus.* Given that these three bacteria are dominant members of the URT community and tend to occur together (42), the reduced diversity in *S. pneumoniae* colonisation state likely reflects their combined dominance of these organisms which suppresses less abundant taxa and reducing diversity.

A key finding in our study is the negative association between *S. pneumoniae* colonisation and both *Granulicatella* and *Filifactor*. This pattern is important because it points to potential avenues for novel interventions aimed at reducing *S. pneumoniae* colonisation given the persistent colonisation prevalence in the post PCV era. Although PCVs have substantially reduced disease, sustaining and extending these gains may require strategies that limits URT colonisation. Within the complex URT ecosystem, numerous microorganisms can interact antagonistically via antimicrobial peptide secretion or nutrient competition (1, 43). When *S. pneumoniae* participates in these interactions, the outcome may influence whether pneumococcal colonisation is sustained or not. For example, bacteriocins produced by the oral streptococcus, *Streptococcus salivarius* has been shown to inhibit the growth of *S. pneumoniae* in vitro studies (43). In our MaAsLin3 analysis, high prevalence of *Granulicatella* and high relative abundance of *Filifactor* were significantly associated with *S. pneumoniae* non-colonisation. Our finding suggests that *Granulicatella* and *Filifactor* may contribute to colonisation resistance. *Kelly MS* et al. also reported that high relative abundance of *Granulicatella* was associated with low *S. pneumoniae* colonisation in children 1–23 months of age in Botswana (44). The mechanism underlying their associations with *S. pneumoniae* is unknown. However, for *Granulicatella, S. aureus* may play a central role. *Granulicatella* lacks the biosynthetic pathways to produce Vitamin B6 (pyridoxal) (45) which is essential for its growth. Consequently, it often requires co-culture with *S. aureus* which releases sufficient vitamin B6 to support its growth in vitro (46). This cross-feeding phenomenon may parallel interactions in the upper respiratory tract where *Granulicatella* could rely on co-habiting bacteria such as *S. aureus* for survival. If *Granulicatella* abundance depends on *S. aureus*, then its negative association with *S. pneumoniae* may indirectly reflect the well-established antagonism between *S. aureus* and *S. pneumoniae*. High *S. aureus* levels in the upper respiratory tract have been reported to occur when *S. pneumoniae* is absent and this state may coincide with high *Granulicatella* prevalence. This remains a hypothetical mechanistic explanation and subsequent research should prioritise shotgun metatranscriptomics along with *in vitro* co-culture and growth inhibition assays to identify the pathways or substances that are responsible for the interactions observed.

The primary strength of this study is in its contribution in describing the upper respiratory tract of healthy older children in Africa, a notably understudied population. Secondly, the longitudinal design of our study helps account for temporal variability often missed in cross-sectional studies. Our study is not without limitation. Notably, the interval between visits is considered long for studying microbiome changes over time meaning that short-term changes were likely missed. Frequency of sampling should be increased in future studies. Secondly, the pause in sample collection between visits four and five due to COVID-19 mitigation measures may have resulted in significant changes in the microbiome, and participants’ environmental exposures during this period were unknown. Finally, our 16S rRNA analysis precluded species-specific insights, and this limited the interpretation of some our observations.

## Conclusion

In summary, this study delineates the URT microbiome of older Gambian children, establishing *Streptococcus* and *Prevotella* as dominant bacteria and describing the influence of environmental factors affecting microbiome composition. We further showed that *Granulicatella* and *Filifactor* were associated with reduced colonisation by *S. pneumoniae* which underscores the need to further investigate the mechanism of this resistance for potential therapeutic exploration.

## List of abbreviations

ASV: Amplicon Sequence Variants
CLR: Centered Log-Ratio
EQA: External Quality Assurance
PCoA: Principal Coordinate Analysis
STGG: Skim milk-Tryptone-Glucose-Glycerol
URT: Upper Respiratory Tract

## Declarations

## Ethics approval and consent to participate

Written informed consent was obtained from the parents or guardians of all participants prior to enrolment in the study. Each parent/guardian received a copy of the participant information sheet and consent form, which was explained in the local language to those not fluent in English. The study was approved by the Joint Medical Research Council Unit The Gambia at the London School of Hygiene and Tropical Medicine and the Government of The Gambia ethics committee. All procedures were performed in accordance with relevant guidelines and regulations.

## Consent for publication

Not applicable

## Availability of data and materials

The sequence data generated in this study have been deposited in the SRA under the BioProject accession PRJNA1538550. The dataset used for the microbiome analyses in this paper is available on Figshare at https://doi.org/10.6084/m9.figshare.34056576. The metadata of the study participants are available upon request, given the requirement for participant confidentiality under the informed consent obtained for this study. Access can be requested through the Gambia Government/MRC Joint Ethics Committee, facilitated by MRC Unit The Gambia (http://www.mrc.gm/) through Martin Antonio, or Dam Khan. All other data are available within the article and supplementary files.

The R code used for the analyses described in this paper is available on GitHub at https://github.com/Bosskhan/Upper-respiratory-microbiome-composition-and-pneumococcal-colonisation-in-The-Gambia, and archived on Zenodo at https://doi.org/10.5281/zenodo.23121184

## Authors’ contributions

BKA, MA, NIM, CLD, and MPN conceived and designed the study. PN, AW and MAK managed sample collection and data tracking. PET, SAT, DK, TM and GJ performed laboratory processing. DK and JLE conducted the data analysis. DK prepared the figures, generated tables and wrote the initial draft of the Manuscript. MPN, CLD, and MA edited and revised the manuscript. All authors participated in the discussion and interpretation of the results.

## Acknowledgements

We express our sincere gratitude to the study participants and their guardians for their invaluable cooperation and time. We are deeply grateful to the Foni community and village heads for the support and facilitation of our fieldwork. Finally, we thank the entire ReMAC laboratory staff for their assistance in sample processing.

