## Supplementary Information for "Upper respiratory tract bacterial microbiome features associated with *Streptococcus pneumoniae* colonisation and environmental factors among healthy older children in a rural setting in The Gambia"

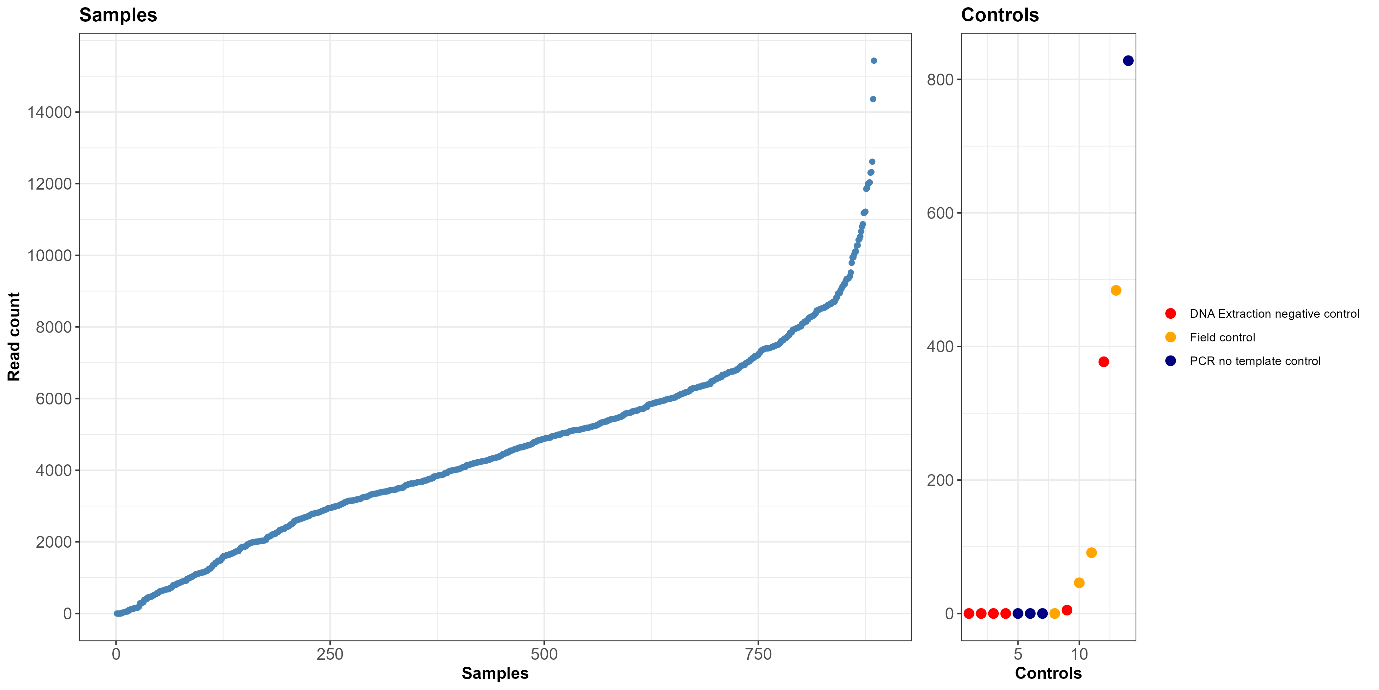

Figure S1: Number of reads in samples and controls. Each dot represents an individual sample or control. Samples and controls are arranged horizontally based on their read count with those having the highest counts towards the right. The y-axis scale differs between the sample and controls plots.

Table S1**:** Number of NP/OP samples processed and retained for 16S rRNA final analysis after exclusion of contaminant and spurious ASVs, and samples with fewer than 1000 reads

| Visit | Swabs processed for both microbiology culture and 16S rRNA sequencing (%) | Swabs retained for 16S rRNA analysis post-QC and sample filtering (%) | Percentage of samples retained per visit |
| --- | --- | --- | --- |
| 1 | 134 (15.1) | 111 (13.9) | 82.8 |
| 2 | 126 (14.2) | 112 (14.0) | 88.9 |
| 3 | 119 (13.4) | 104 (13.0) | 87.4 |
| 4 | 114 (12.9) | 98 (12.3) | 86.0 |
| 5 | 106 (12.0) | 102 (12.8) | 96.2 |
| 6 | 98 (11.1) | 91 (11.4) | 92.9 |
| 7 | 97 (11.0) | 94 (11.8) | 96.9 |
| 8 | 91(10.3) | 87 (10.9) | 95.6 |
| Total | 885 (100) | 799 (100) | 90.3 |

Table S2: Distribution of participants by number of samples retained in the final 16S rRNA dataset across all eight visits following participant withdrwals and all quality control checks

| Number of samples included in final dataset | number of participants | % |
| --- | --- | --- |
| 0 | 1 | 0.7 |
| 1 | 5 | 3.7 |
| 2 | 9 | 6.7 |
| 3 | 11 | 8.2 |
| 4 | 7 | 5.2 |
| 5 | 8 | 6.0 |
| 6 | 18 | 13.4 |
| 7 | 33 | 24.6 |
| 8 | 42 | 31.3 |
| Total | 134 | 100 |

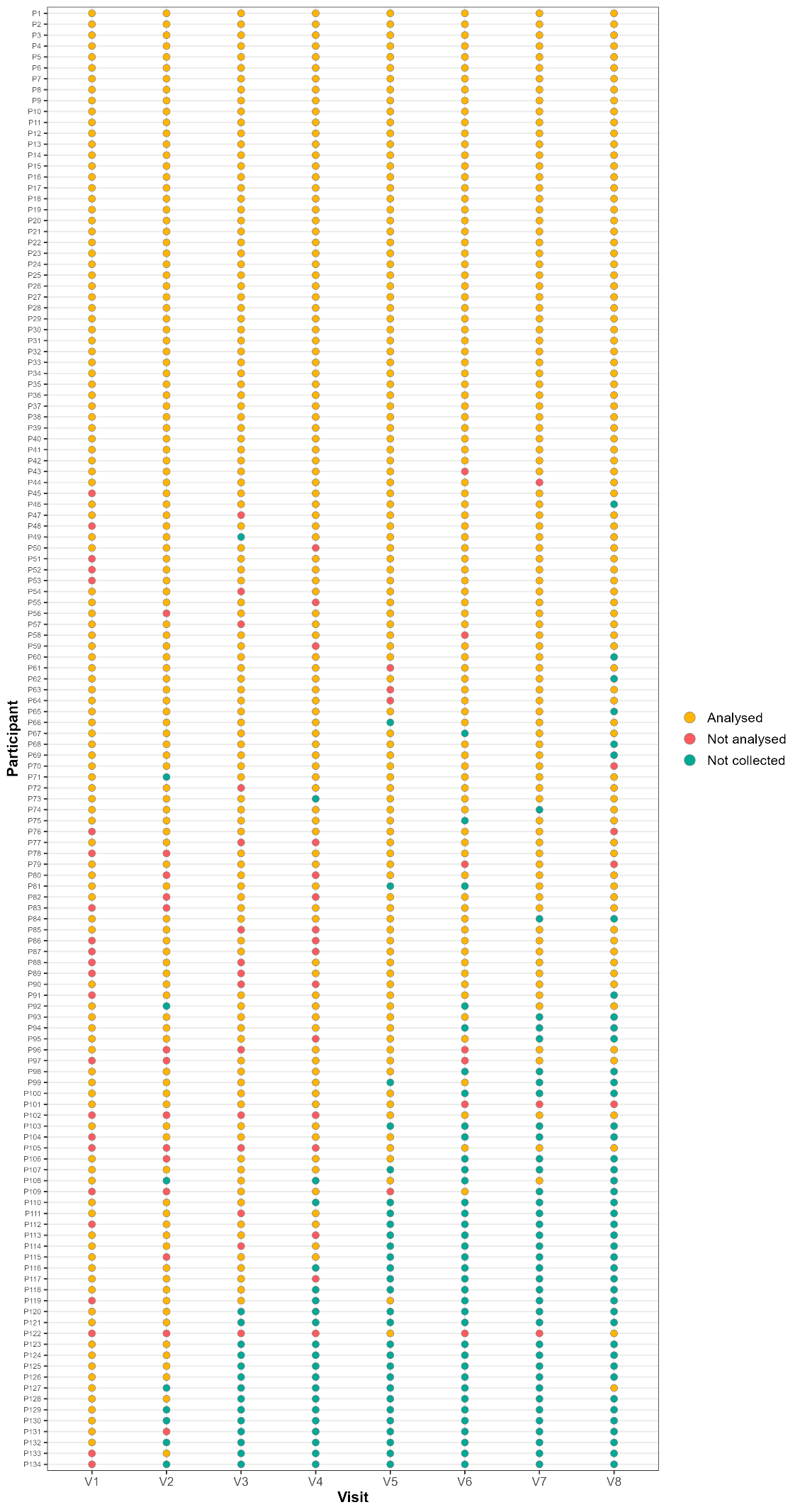

Figure S2: Number of samples included in the final 16S rRNA gene analysis for each participant.

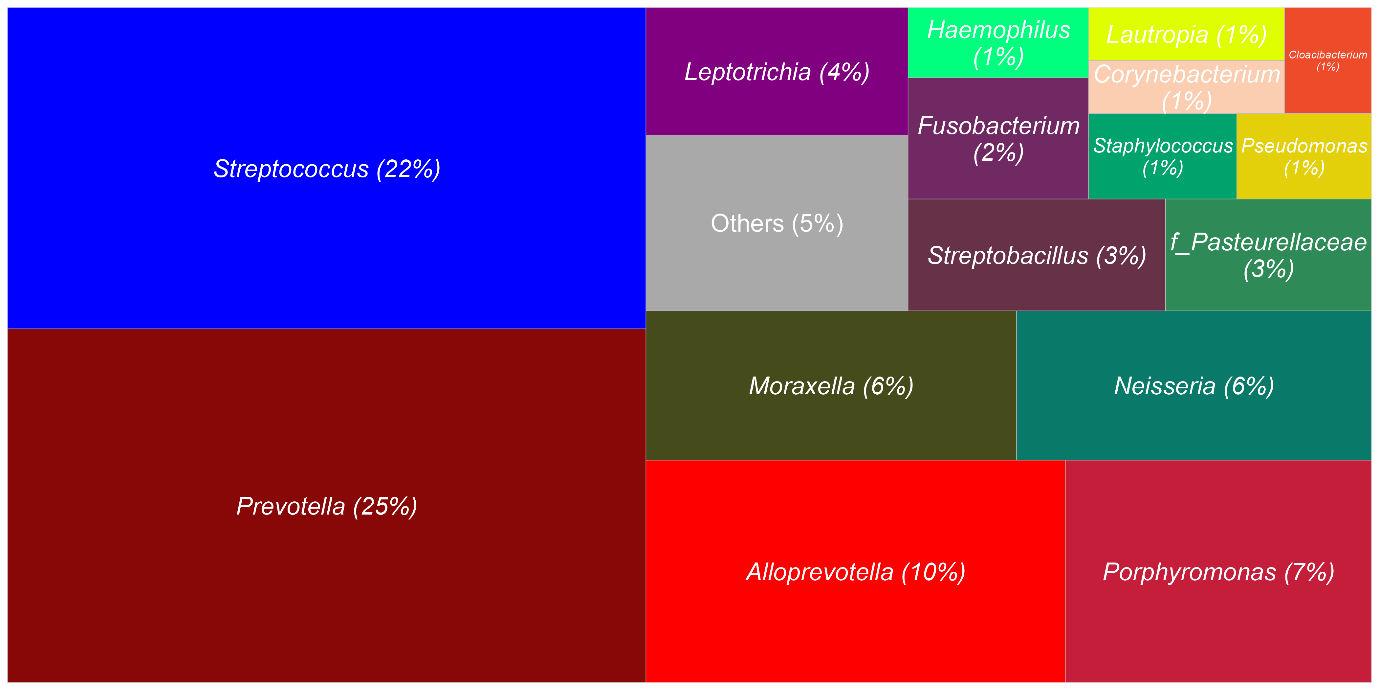

Figure S3: The treemap illustrates the distribution of dominant genera across all samples. The size of each segment corresponds to the percentage of samples dominated by that specific genus. Genera dominating less than 1% of samples have been grouped into ‘Others’

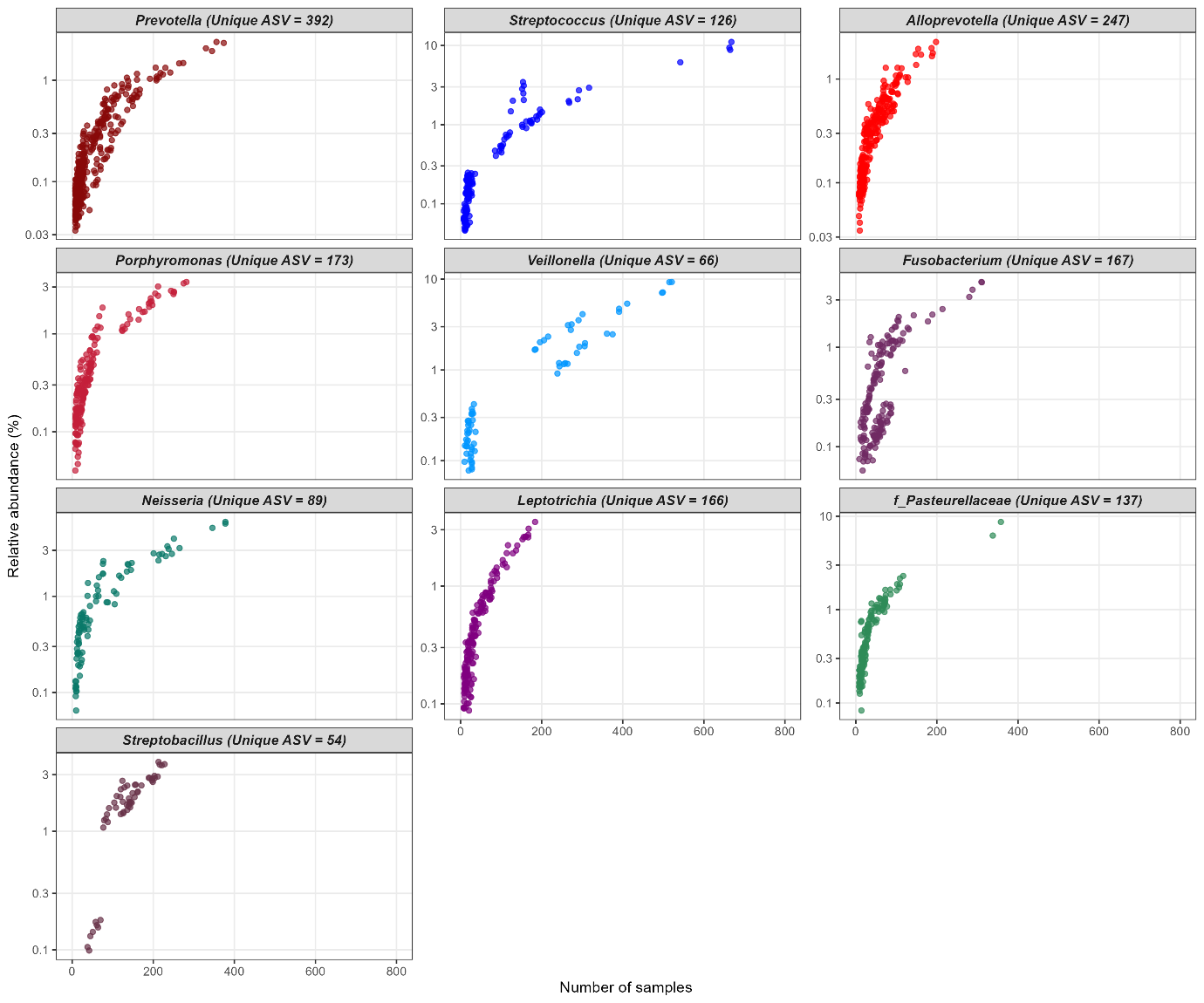

Figure S4: Count of unique ASVs across all samples for the top 10 genera. The relative abundances of each ASV within their respective genera is shown on the Y-axis. The Y-axis scale differs for the individual plots.

Table S2: Association between NP/OP Shannon diversity and environmental and sociodemographic factors

| Covariate | Estimate | SE | P value |
| --- | --- | --- | --- |
| Sex (Male Vs Female) | -0.073 | 0.06 | 0.228 |
| Age at visit (per year) | 0.029 | 0.013 | **0.019*** |
| Number of siblings (per sibling) | -0.02 | 0.02 | 0.218 |
| Season (Wet Vs Dry) | 0.073 | 0.045 | 0.105 |
| Presence of pets in the compound (Yes Vs No) | -0.063 | 0.052 | 0.229 |
| Presence of livestock in the compound (Yes Vs No) | -0.012 | 0.06 | 0.847 |
| Source of fuel (Firewood Vs Charcoal) | 0.074 | 0.06 | 0.186 |
| Waste disposal method (Open burning in the household Vs Open burning outside the household) | 0.031 | 0.122 | 0.798 |

*Statistically significant

Table S3: Differential abundance and prevalence of URT taxa associated with household environmental and sociodemographic factors (q < 0.1)

| Covariate | Model | Taxon | Enrichment group | Coefficient | Std. Error | q-value |
| --- | --- | --- | --- | --- | --- | --- |
| Sex | Abundance | *Campylobacter* | Male | 0.219 | 0.080 | 0.071 |
| Presence of pets | Prevalence | *Dolosigranulum* | No pets | 0.707 | 0.242 | 0.024 |
|  | Prevalence | *Gemella* | No pets | 0.828 | 0.302 | 0.039 |
|  | Abundance | *Megasphaera* | Pets present | 0.245 | 0.084 | 0.042 |
| Presence of livestock | Abundance | *Campylobacter* | Livestock present | 0.273 | 0.09 | 0.063 |
|  | Prevalence | *Lactococcus* | Livestock present | 0.933 | 0.356 | 0.049 |
| Source of fuel | Abundance | *Lactococcus* | Charcoal users | 0.631 | 0.214 | 0.035 |
|  | Prevalence | *Catonella* | Firewood users | 0.808 | 0.233 | 0.005 |
|  | Prevalence | *f_Anaerovoracaceae* | Firewood users | 0.608 | 0.233 | 0.056 |
| Waste burning location | Prevalence | *Pseudomonas* | Burned inside | 0.944 | 0.395 | 0.096 |
|  | Prevalence | *f_Lachnospiraceae* | Burned inside | 0.671 | 0.262 | 0.072 |
|  | Prevalence | *Treponema* | Burned outside | 0.880 | 0.274 | 0.012 |
